# Validation of a high-throughput fluorescent Capillary Electrophoresis Sodium Dodecyl Sulfate method for monoclonal antibody size heterogeneity assessment

**DOI:** 10.64898/2026.07.15.738750

**Authors:** Kyle D Luttgeharm, Manu Grover, Sheng-Yuan Huang, Whitney A Pike

**Affiliations:** Agilent Technologies, 2450 SE Oak Tree Ct, Ankeny, Iowa, USA; Agilent Technologies, 5301 Stevens Creek Blvd, Santa Clara, California, USA

**Author notes:** Corresponding Author: Kyle D Luttgeharm. **Competing Interest:** KDL, MG, SYH, and WAP all declare employment and equity or stock holdings with Agilent Technologies.

**Keywords:** Capillary Electrophoresis SDS, Monoclonal Antibody, Antibody Critical Quality Attributes, Antibody Method Validation

## Abstract

Fluorescent capillary gel electrophoresis (CGE) with sodium dodecyl sulfate (CE-SDS) provides a powerful, high-sensitivity alternative to ultraviolet (UV)-based detection for characterizing therapeutic monoclonal antibodies (mAb). Regulatory and standards organizations, such as the United States Pharmacopeia (USP), include only UV based CE-SDS methods, hindering adoption of alternative detection methods. There is growing opportunity to expand beyond exclusively UV-based CE-SDS methods. In this study, we present an analytical validation of a light-emitting diode (LED) fluorescence-based parallel CE-SDS method for both non-reduced and reduced analysis of therapeutic antibodies. Using the NISTmAb reference material as a model system, size heterogeneity critical quality attributes (CQAs) including monomeric purity (non-reduced conditions), percent glycosylation (reduced conditions), and percent thioether (reduced conditions) were assessed. The fluorescence method demonstrated high specificity and precision with relative standard deviation (RSD) values <1% for monomeric purity and glycosylation, and <3% for thioether, as well as robust performance across variations in injection voltage, electrophoresis voltage, labeling temperature, and Labeling Buffer concentration. Ruggedness testing across users and reagent lots confirmed reproducibility, and accuracy assessments showed strong agreement with reported values from the National Institute of Standards (NIST) and traditional UV detection measurements. Linearity studies yielded coefficient of determination (R^2^) values >0.99 for both non-reduced and reduced analyses. These results highlight the high sensitivity, stable baseline performance, and suitability of LED fluorescence-based parallel CE-SDS as a validated, higher-throughput alternative to traditional UV-based methods for mAb quality control (QC).

## Introduction

Therapeutic monoclonal antibodies (mAb) target specific antigens, making them ideal therapeutics for complex diseases such as autoimmune disorders, asthma, and cancers. As of January 2026, 146 therapeutic mAbs have been approved for use in the United States with an average of ∼10 new approvals each year since 2015 [1], indicating increased adoption of biologic therapeutics. The majority of these therapeutic mAbs are built upon the immunoglobulin G (IgG) backbone and consist of two heavy chains (H, ∼50 kDa) and two light chains (L, ∼25 kDa) held together by inter-chain disulfide bonds. The mAb can be further modified by the addition of post-translational modifications, such as glycosylation, which contribute to the overall biophysical stability [2]. Therapeutic mAbs are mainly produced in recombinant mammalian cell lines, as these cells can carry out essential post-translational modifications.

During expression of any biological product, impurities derived from both the cellular production system and purification process are generated. For therapeutic mAbs, these impurities include antibody fragments, host cell proteins, and non-glycosylated mAb species [2]. Development and manufacturing of therapeutic mAbs therefore require strict quality control (QC) analysis for full characterization of the therapeutic and to ensure batch-to-batch consistency. QC analysis of therapeutic mAbs includes glycosylation profiling, along with both charge and size heterogeneity. The size heterogeneity Critical Quality Attributes (CQAs) analyzed include monomeric purity, percent glycosylation, and percent non-reducible/thioether species [3]. Monomeric purity consists of the percent monomer compared to the total when analyzed under non-reduced conditions, [4]while percent glycosylation is the relative amount of glycosylated heavy chain compared to the total of the non-glycosylated and glycosylated heavy chain [4] when analyzed under reduced conditions. Non-reducible/thioether species result from the elimination of one sulfur atom from the disulfide bond [5] and have been shown to form between a light and heavy chain [6]. Percent thioether is determined by comparing the non-reducible/thioether species to the total when analyzed under reduced conditions [4].

Capillary gel electrophoresis (CGE) in the presence of sodium dodecyl sulfate (CE-SDS) has emerged as a primary tool for size heterogeneity assessment over traditional SDS-poly acrylamide gel electrophoresis (PAGE) due to its high-resolution separations, quantitative reproducibility, and the availability of automated systems [7]. Methods for the analysis of mAbs by CE-SDS using UV absorption for protein detection have been published by organizations such as the United States Pharmacopeia (USP) and are widely used for the QC of mAbs [8, 9]. While detection by UV absorption allows label-free analysis, capillary and lamp temperature variations often result in baseline instability [10-12]. These instabilities lead to variability in peak areas and difficulty detecting low concentration species, thus impacting the CQA measurements of the mAb [11, 13, 14]. In contrast to UV absorption, fluorescence-based detection using either laser-induced or light-emitting diode (LED)-induced excitation generates more stable baselines, lower background noise, and superior signal intensity, which allows for detection of low-concentration impurities with greater precision [15-18]. Both laser- and LED-induced fluorescence result in a more stable baseline compared to UV [16]. Additionally, LED-based fluorescence has been shown to have improved baseline stability compared to laser-induced fluorescence [15]. Despite the potential advantages of fluorescence-based CE-SDS including baseline stability and sensitivity, it has not been widely adopted for therapeutic mAb QC. The slower adoption of fluorescence detection may reflect its absence from regulatory compendia, along with the long-standing acceptance of UV-based methods.

Traditionally, CE-SDS has been performed on single capillary systems using time corrected peak areas for size heterogeneity CQA analysis. These single capillary systems have typical electrophoresis times of around 20 min per sample and can run up to 96 samples without user intervention. With the increasing demand for therapeutic mAbs, higher throughput analysis is required. Parallel capillary electrophoresis systems allow for multiple samples to be analyzed simultaneously, vastly increasing throughput compared to traditional single capillary systems. For instance, the system used in this study can analyze 12 samples in about 30 min with up to 288 samples analyzed without user intervention. While parallel CE-SDS systems allow for increased throughput, it also increases variation in migration time and corrected peak areas due to differences in capillary length, diameter, and window location [19]. Internal standards can be used to mitigate these sources of variation, allowing for corrections to sample migration time, calculation of sample size, and the use of peak area ratios for relative quantification [20]. The use of internal standards for size and concentration calculations has not been previously included in methods such as those published by USP [8], further demonstrating the need for additional validated methods.

Here, we present data for the validation of an LED fluorescence-based parallel CE-SDS method using the Agilent ProteoAnalyzer system. The ProteoAnalyzer system uses constant LED fluorescence with detection by a charge-coupled device (CCD) camera. A 12-channel parallel capillary array allows for analysis of 12 samples simultaneously. The NISTmAb, Humanized IgG1_*K*_ Monoclonal Antibody, was analyzed under non-reduced conditions to determine monomeric purity, intact mAb size, and intact mAb relative concentration. Reduced conditions were used to determine percent glycosylation and percent thioether, along with Light Chain (L), Non-Glycosylated Heavy Chain (NGH), Heavy Chain (H), and thioether size and relative concentration. RSDs are reported for all values as the measure of precision. Specific tests were run to validate assay specificity, precision (instrument and method), robustness (injection, instrument, denaturing/labeling temperature, and 10x Labeling Buffer concentration), ruggedness, accuracy, and linearity. The highly reproducible results provide a validated method for mAb analysis using parallel CE-SDS using LED fluorescence.

## Material & Methods

Validation experiments followed the general guidelines outlined by the International Council for Harmonisation of Technical Requirements for Pharmaceuticals for Human Use (ICH) in Validation of Analytical Procedures Q2(R2) [21]. The NISTmAb, Humanized IgG1_*K*_ Monoclonal Antibody (Reference Material 8671, Lot 14HB-D-002) was analyzed as the representative test sample. Monomeric purity, percent glycosylation, and percent thioether were chosen as the key CQAs due to their inclusion by NIST in the Reference Material Information Sheet [22].

All fluorescent analysis was completed on the Agilent ProteoAnalyzer system (Agilent p/n M5350-64000) using the Agilent ProteoAnalyzer 12-Capillary Array (Agilent p/n M5350-64001), and the Agilent Broad Range P240 reagent kit (Agilent p/n 5191-6642) per the manufacturer’s instructions. The USP Monoclonal IgG System Suitability RS (USP p/n 1445550) was run as the first and last set of samples, and was required to pass the system suitability requirements described in USP <129> [8]. All samples were diluted in 1x phosphate-buffered saline (PBS) (Sigma p/n D8537). Samples were prepared at 1,500 ng/µL for all injections except for the linearity study where samples were prepared at 1,000 ng/µL, 1,200 ng/µL, 1,500 ng/µL, 1,800 ng/µL, and 2,000 ng/µL. All concentrations were verified with a Thermo Fisher Scientific NanoDrop One using the Protein 280 analysis mode and an E1% extinction coefficient of 14.2.

### LED fluorescent-based CE-SDS sample preparation

Unless otherwise stated, the manufacturer’s sample preparation protocol was used. For non-reduced analysis, 45 µL 10x Labeling Buffer, 45 µL 250 mM iodoacetamide, 3.9 µL Protein Fluorescent Dye, and 340 µL CE-grade water were mixed to prepare the Reaction mix. For reduced analysis, the Reaction mix was composed of 45 µL 10x Labeling Buffer, 4.5 µL 1 M Dithiothreitol (DTT), 3.9 µL Protein Fluorescent Dye, and 381 µL CE-grade water. Samples were diluted 1:29 with the appropriate Reaction mix to make master mixes. 30 µL of the master mix was aliquoted into wells of an Eppendorf Twin.Tec semi-skirted 96-well plate (Eppendorf p/n 951020303). Blanks were prepared by adding 1x PBS in lieu of sample. The prepared plates were sealed and heated at 70 °C for 10 min on a thermocycler to denature and covalently label the samples with the fluorescent dye.

For method precision experiments, the NISTmAb was prepared by mixing 1 µL sample with 29 µL Reaction mix directly in the 96-well plate. Samples were mixed using a MicroPlate Genie (Scientific Industries) for 2 min at 3,000 rpm. For the labeling temperature robustness study, samples were heated at 68 °C, 70 °C, and 72 °C. For the Labeling Buffer volume robustness study, Reaction mixes were prepared with 44 µL, 45 µL, and 46 µL 10x Labeling Buffer, while all other Reaction mix reagent volumes remained the same as previously described.

### LED fluorescent-based CE-SDS instrument operation

The ProteoAnalyzer system was prepared according to the manufacturer’s instructions. Briefly, 1 mL 1x Protein Inlet Buffer, 1 mL sub-micron filtered distilled water, and 1 mL Agilent Capillary Storage Solution (Agilent p/n GP-440) were added to the wells of rows A, B, and H, respectively, of a Fisher 1 mL DeepWell 96-well plate (Fisher Scientific p/n 12-566-120) and placed on the ProteoAnalyzer in drawer “B”. The inlet buffer and CE-grade water were removed and replaced prior to the first run of the day. Broad Range Protein gel, Protein Conditioning Solution, and 1 M NaOH were placed on the appropriate instrument lines. Prior to the first run of the day, the “Daily Conditioning Flush” was completed. Samples were analyzed using the *5191-6642LM – ProteoAnalyzer Broad Range Kit LM-only* method. For reduced samples, the default sample injection of 7 kV for 10 s was used. For non-reduced samples the injection was decreased to 7 kV for 6 s per the manufacturer’s recommendation [23]. Electrophoresis was completed at 9 kV for 20 min.

For instrument injection robustness, samples were analyzed using injection voltages of 6.9 kV, 7.0 kV, and 7.1 kV with a 10-s injection time for reduced samples and 6-s injection time for non-reduced samples. For instrument separation robustness, samples were analyzed at electrophoresis voltages of 8.9 kV, 9 kV, and 9.1 kV, all with 20-min separations.

### LED fluorescent-based CE-SDS data analysis

Runs were analyzed using the Agilent ProSize data analysis software v5.0.1.6. All samples were analyzed with the same ProSize data analysis software settings. Non-reduced analysis settings were a peak width of 4 s, 20 RFU minimum peak height, and the Valley-to-Valley selected using one extra valley point. Analysis settings for reduced samples were a peak width of 1 s, 30 RFU minimum peak height, with the Valley-to-Valley selected using one extra valley point. For non-reduced samples, protein sizes were calculated using the NISTmAb and its impurities as the ladder (L 23 kDa, H 50 kDa, HL 73 kDa, H:H 101 kDa, H:H:L 124 kDa, intact monomer 148 kDa) as previously described [24, 25] to correct for sample migration rate differences resulting from non-reduced disulfides when compared to a linear polypeptide ladder [26, 27]. For reduced samples, the Agilent P240 Broad Range ladder was used.

Relative concentration (ng/µL) is automatically calculated by normalizing the sample-corrected peak areas with that of the 6 kDa Lower Marker (LM) supplied in the Agilent 10x Labeling Buffer. This normalization adjusts for capillary-to-capillary variation, as previously described [15, 20]. Non-reduced monomeric purity values were determined using the relative concentration of the intact mAb divided by the relative concentration of all detected samples. Percent glycosylation and percent thioether were determined using previously established equations [4] from reduced analysis.

### UV absorbance based CE-SDS sample preparation

The reduced and non-reduced NISTmAb samples were prepared according to the manufacturer’s protocol for the SCIEX BioPhase 8800 system. The reduced NISTmAb was diluted with SDS-MW sample buffer (SCIEX, p/n A30341) to a concentration of 1 mg/mL. Next, 2 µL of 10 kDa internal standard (SCIEX, p/n A26487) and 5 µL of 2-mercaptoethanol were added to 95 uL of the diluted sample, mixed thoroughly, spun down and heated at 70 °C for 10 min. The same process was followed for the non-reduced NISTmAb, but with 2 µL of 10 kDa internal standard and 5 µl of 250 mM iodoacetamide. The samples were mixed thoroughly, spun down and heated at 70 °C for 10 min. After the samples cooled down to room temperature, 100 µL of each mixture was added to the BioPhase 8800 sample inlet plate (SCIEX, p/n # 5080313), and centrifuged at 30 g for 4 min.

### UV absorbance based CE-SDS instrument operation

The SCIEX BioPhase 8800 system was prepared according to the manufacturer’s instructions. Reagents included: CE grade water (SCIEX, p/n C48034), 0.1 N HCl, 0.1 N NaOH and CE-SDS gel buffer (SCIEX, p/n A30341). All samples were analyzed using the 8 bare fused silica capillary cartridge (SCIEX, p/n 5080121), with the Reduced CE-SDS Separation and Non-Reduced CE-SDS Separation methods. Electrokinetic injection was performed at 5 kV for 30 s. Separation of reduced and non-reduced samples was 15 kV for 40 and 50 min, respectively. A CE-SDS Conditioning method was performed at the beginning of each day and a CE-SDS Shutdown method at the conclusion of the day.

### UV absorbance based CE-SDS data analysis

The results were analyzed using the SCIEX BioPhase data analysis software v1.4.100. For the identification and analysis parameters of reduced NISTmAb peaks, the width of the moving filter was set at 50 s, the positive threshold at 0.5%, minimum cluster distance at 1, and the suspend integration ranging from 0 to 13 min. For the non-reduced samples, the width of the moving filter was set at 50 s, the positive threshold at 0.03%, minimum cluster distance at 50, and the suspend integration ranging from 0 to 14.7 min.

### Baseline drift analysis

Four replicate injections of PBS blank were acquired on four capillaries of the BioPhase 8800 cartridge, and four replicate injections of PBS blanks on five capillaries of the ProteoAnalyzer. The analysis window was set to 800–2,400 s on the BioPhase 8800 and to 0–1,200 s on the ProteoAnalyzer, with the 470–650 s interval excluded due to the presence of the system peak and Lower Marker. The selected times on both systems represents the complete electrophoresis time. To characterize the baseline drift, the analysis window was divided into 30 s segments and each segment was fitted by linear regression, following Chapter 9 (Noise determination) of the Agilent OpenLab CDS Data Analysis Reference Guide with slight modifications [28]. In brief, the residual standard deviation was calculated according to equation 1, where *nn*_*ii*_ is the measured signal, *nn*_*ii*_ is the corresponding value on the fitted line, and *n* is the total number of points in the segment. The linear fit was calculated for each segment and the median value was taken as the noise for the drift (σσ). To avoid overestimating the short-term noise by including long-term drift, *RR*_*ssssss*_ was determined with a 30 s window and is denoted σσ. The signals were then fitted by linear regression over time intervals of 1,600 s from the start of the analysis window on the BioPhase 8800, and over a fixed interval of 10,199 s on the ProteoAnalyzer. The resulting slopes were used to calculate the net drift (Δ) over each interval, and the drift-to-noise ratio (% Drift /σσ,) was used to compare the two systems: 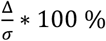, with the unit of mAU/s for Biophase 8800 and RFUs/s for ProteoAnalyzer, respectively.

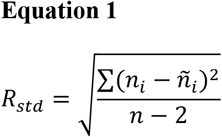

### Fluorescent limit of detection and limit of quantification

To evaluate the limit of detection (LOD) and limit of quantification (LOQ), a serial dilution of non-reduced NIST mAb was prepared from 50 ng/µL to 1.5 ng/µL, representing the bottom end of the manufacturer’s linear dynamic range of 2 – 2,000 ng/µL of Bovine Serum Albumin. The samples were prepared in triplicate and analyzed as previously described. Each replicate was used to create a unique calibration curve. As in the linearity experiments, the non-reduced monomer peak in the electropherogram was used as the main peak for quantification. For each measurement, the noise was determined over a baseline window centered on the mean migration time of the monomeric peak and spanning ± 3 × the full width at half maximum (FWHM) of that peak. The LOD and LOQ were obtained by interpolating a signal equal to three and ten times this noise onto the calibration curve, as previously described [29]. The LOD and LOQ values for each of the three replicates were averaged.

## Results

### Specificity

Specificity of an analytical method can be evaluated by demonstrating the absence of interfering signal when compared to a blank [21, 30]. Here, specificity was evaluated by comparing capillaries containing sample to those containing blank matrix using both the fluorescent CE method and traditional UV detection (Figure 1 and Figure S1, S2). For the fluorescent method, the blank analysis under non-reducing and reducing conditions clearly showed a system peak resulting from unconjugated fluorescent dye eluting first, followed by the 6 kDa LM used for sample alignment and relative quantity, with a small impurity eluting shortly after the LM. This impurity is presumably from the LM and was excluded from all calculations. The baseline after the LM impurity did not contain peaks that fit the integration parameters, indicating a lack of interfering signals (Figure 1a and b, Figure S1). Both the NISTmAb non-reduced and reduced CE-SDS profiles (Figure 1a and c) were consistent with those previously published by NIST [4, 31]. In contrast the baseline of the UV method (Figure 1c and d, Figure S2) showed significant variation, particularly under non-reduced conditions (Figure 1c, Figure S2).

**Fig. 1.**
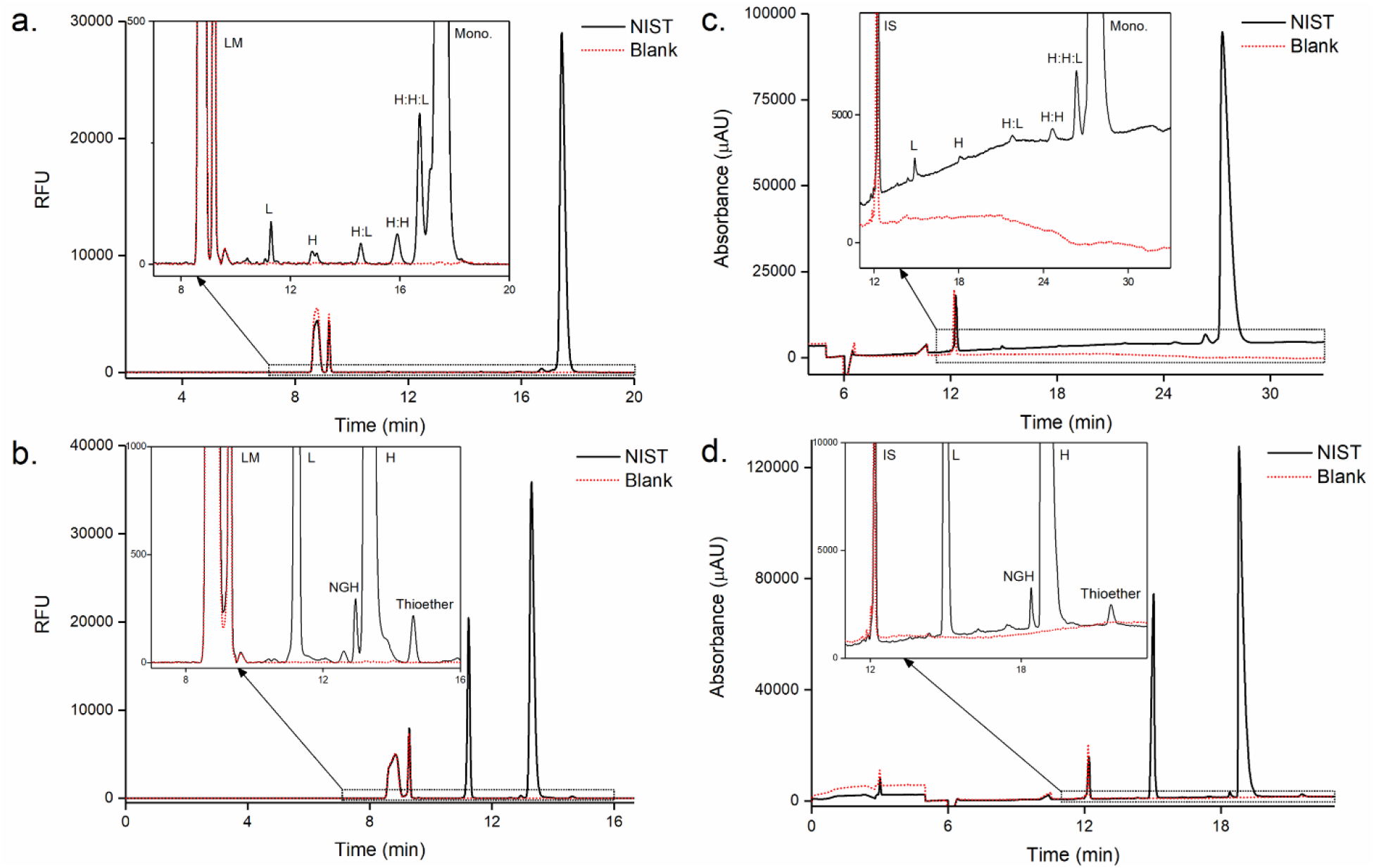
Representative electropherograms of the non-reduced and reduced NISTmAb analyzed using the LED-induced fluorescence-based method (a and b, respectively) and the UV-based method (c and d, respectively). Samples traces (black) are overlaid with a blank to demonstrate the baseline stability (red). LM = Lower Marker, IS = Internal Standard, L = Light Chain, NGH = Non-glycosylated Heavy Chain, H = Heavy Chain, Mono = Monomer.

To compare the baseline behavior of the two detection systems, we treated the long-term drift and short-term noise separately. The noise was extracted by dividing the analysis window into segments of fixed length and calculating the residual standard deviation (*R*_*std*_) of the local linear fit. The drift-to-noise ratio (% Drift /σσ) was used to compare baseline stability between the two systems. After excluding the lower marker region from the fluorescence electropherogram, the drift measured on the fluorescent detector over a 1019 s interval and compared with that measured on the UV detector over a 1,600 s interval. Both represent the complete electrophoresis time excluding any system and marker peaks. The average % Drift /σσ was 0.03 ± 0.02 % for the FL detector, compared with 7.59 ± 3.79 % for the UV detector (Figure S3) demonstrating that the LED based fluorescence detector on the ProteoAnalyzer maintains a more stable baseline than the UV detector on the BioPhase 8800. The clean, stable, baseline of the blank sample and well-defined sample peaks demonstrate the specificity of the fluorescent CE-SDS assay for mAb analysis. The fluctuating UV baseline increased the complexity of peak integration compared to the fluorescent method [14].

### Precision

ICH Q2(R2) guidelines state that precision may be determined by measuring a minimum of 6 samples at 100% of the test concentration [21]. Instrument precision was determined using a master mix containing NISTmAb and Reaction mix at a 1:29 ratio and aliquoted into wells 1 – 6. The monomeric purity (Table 1), relative concentration, and size (Table S1) of the non-reduced NISTmAb was determined. All values showed high precision with RSD values of 0.053%, 0.831%, and 1.311%, respectively.

**Table 1.** Average monomeric purity, %glycosylation, %thioether, and RSD values for the NISTmAb from six independent capillaries used to evaluate instrument precision of the LED fluorescence-based parallel CE method. N = 6.

|  | Average | Precision (RSD) |
| --- | --- | --- |
| Monomeric Purity<sup>(a)</sup> | 98.07% | 0.053% |
| %Glycosylation<sup>(b)</sup> | 99.38% | 0.011% |
| %Thioether<sup>(b)</sup> | 0.40% | 2.534% |

<sup>(a)</sup>Measured under non-reducing conditions
<sup>(b)</sup>Measured under reducing conditions

Instrument precision for reduced samples was determined in the same manner as the non-reduced samples, with the size and relative concentration calculated for the L, NGH, H, and thioether species (Table S1), along with the CQAs of percent glycosylation and percent thioether (Table 1). RSD values for the percent glycosylation and percent thioether CQAs were 0.011% and 2.534%, respectively.

To validate the method precision, samples were individually mixed 1:29 (sample : Reaction mix) in wells 1 – 6. The same CQAs as the instrument precision tests were calculated. Slightly higher precision values for relative concentration and size were observed with RSD values of 3.793% and 1.511%, respectively (Table S2). Observed monomeric purity precision value was consistent with that of the instrument precision test. Method precision for reduced samples showed comparable results as the non-reduced analysis with RSD values for percent glycosylation and percent thioether of 0.017% and 0.729%, respectively (Table 2). Relative concentrations and sizes showed slightly higher RSD values than the instrument precision test (Table S2). Slightly higher values for the relative concentrations and sizes were expected due to the inherent nature of individually mixing samples instead of aliquoting from a master mix.

**Table 2.** Average monomeric purity, %glycosylation, %thioether, and RSD values for the NISTmAb from samples individually mixed 1:29 (sample : Reaction mix) used to evaluate the method precision. N = 6.

|  | Average | Precision (RSD) |
| --- | --- | --- |
| Monomeric Purity<sup>(a)</sup> | 98.13% | 0.053% |
| %Glycosylation<sup>(b)</sup> | 99.38% | 0.017% |
| %Thioether<sup>(b)</sup> | 0.41% | 0.729% |

<sup>(a)</sup>Measured under non-reducing conditions
<sup>(b)</sup>Measured under reducing conditions

The consistent CQA values found in both the instrument and method precision tests demonstrate the high level of reproducibility LED fluorescence-based CE-SDS provides for analysis of monoclonal antibodies.

### Robustness

ICH guidelines state that Robustness testing should demonstrate the reliability of an analytical procedure in response to deliberate variations in the analytical procedure parameters [21]. Here, method robustness was assessed with respect to injection voltage, electrophoresis voltage, labeling temperature, and amount of 10x Labeling Buffer. Each test was performed with three replicates per run. The runs were performed at default conditions followed by parameters above and below the default conditions. All three test conditions were averaged and the RSD value was determined from the pooled data.

Injection robustness was evaluated using the default 7 kV injection followed by reinjection of the same wells at both 6.9 kV and 7.1 kV injections (Tables 3 and S3). For electrophoresis robustness, samples were analyzed using the default 9 kV separation followed by reanalysis of the same samples using 8.9 kV and 9.1 kV electrophoresis voltages (Table 3 and S3). Labeling reaction/denaturing temperature robustness was examined by completing the incubation at the initial recommended temperature of 70 °C followed by incubations at 68 °C and 72 °C (Table 3 and S3). Robustness with respect to the amount of 10x Labeling Buffer was conducted by analyzing samples at the default volume of 45 µL, along with samples prepared with 44 µL and 46 µL of 10x Labeling Buffer (Table 3 and S3). Each parameter investigated gave similar results with low RSD values demonstrating the robustness of LED fluorescence-based CE-SDS for antibody analysis.

**Table 3.** Average monomeric purity, %glycosylation, %thioether, and RSD values for the NISTmAb used to evaluate the method robustness. N = 9.

|  |  | Average | Precision (RSD) |
| --- | --- | --- | --- |
| Injection Voltage | Monomeric Purity<sup>(a)</sup> | 98.09% | 0.032% |
|  | %Glycosylation<sup>(b)</sup> | 99.37% | 0.015% |
|  | %Thioether<sup>(b)</sup> | 0.40% | 1.787% |
| Electrophoresis Voltage | Monomeric Purity<sup>(a)</sup> | 98.06% | 0.051% |
|  | %Glycosylation<sup>(b)</sup> | 99.37% | 0.15% |
|  | %Thioether<sup>(b)</sup> | 0.40% | 1.153% |
| Labeling Temperature | Monomeric Purity<sup>(a)</sup> | 98.07% | 0.140% |
|  | %Glycosylation<sup>(b)</sup> | 99.36% | 0.037% |
|  | %Thioether<sup>(b)</sup> | 0.40% | 3.867% |
| Labeling Buffer Volume | Monomeric Purity<sup>(a)</sup> | 98.09% | 0.032% |
|  | %Glycosylation<sup>(b)</sup> | 99.37% | 0.038% |
|  | %Thioether<sup>(b)</sup> | 0.40% | 2.839% |
| <sup>(a)</sup>Measured under non-reducing conditions |  |  |  |
| <sup>(b)</sup>Measured under reducing conditions |  |  |  |

### Ruggedness (intermediate precision)

Per ICH Q2(2) guidelines, ruggedness or intermediate precision should establish the effects of random events on the precision of the analytical procedure. This can include different days, analysts, and reagents [21]. The ruggedness of the fluorescent assay was evaluated with respect to users/days and reagent lots. Two different users analyzed six replicates using the same lots of reagents and instrument under both non-reduced and reduced conditions on different days (Tables 4 and S4). Consistent CQA values between users and days demonstrate the stability of the assay.

**Table 4.** Average monomeric purity, %glycosylation, %thioether, and RSD values for the NISTmAb from two different users on two different days using the same lot of reagents used to evaluate the method ruggedness. N = 6.

|  |  | Average | Precision (RSD) |
| --- | --- | --- | --- |
| User 1 | Monomeric Purity<sup>(a)</sup> | 98.05% | 0.056% |
|  | %Glycosylation<sup>(b)</sup> | 99.35% | 0.012% |
|  | %Thioether<sup>(b)</sup> | 0.40% | 3.226% |
| User 2 | Monomeric Purity<sup>(a)</sup> | 98.08% | 0.042% |
|  | %Glycosylation<sup>(b)</sup> | 99.36% | 0.011% |
|  | %Thioether<sup>(b)</sup> | 0.39% | 2.161% |
| <sup>(a)</sup>Measured under non-reducing conditions |  |  |  |
| <sup>(b)</sup>Measured under reducing conditions |  |  |  |

To evaluate the fluorescent assay’s ruggedness with respect to reagent lots, a second set of reagents was used. Consistent CQA values were found between test cases, further highlighting the ruggedness of fluorescent based CE-SDS (Tables 5 and S5).

**Table 5.**
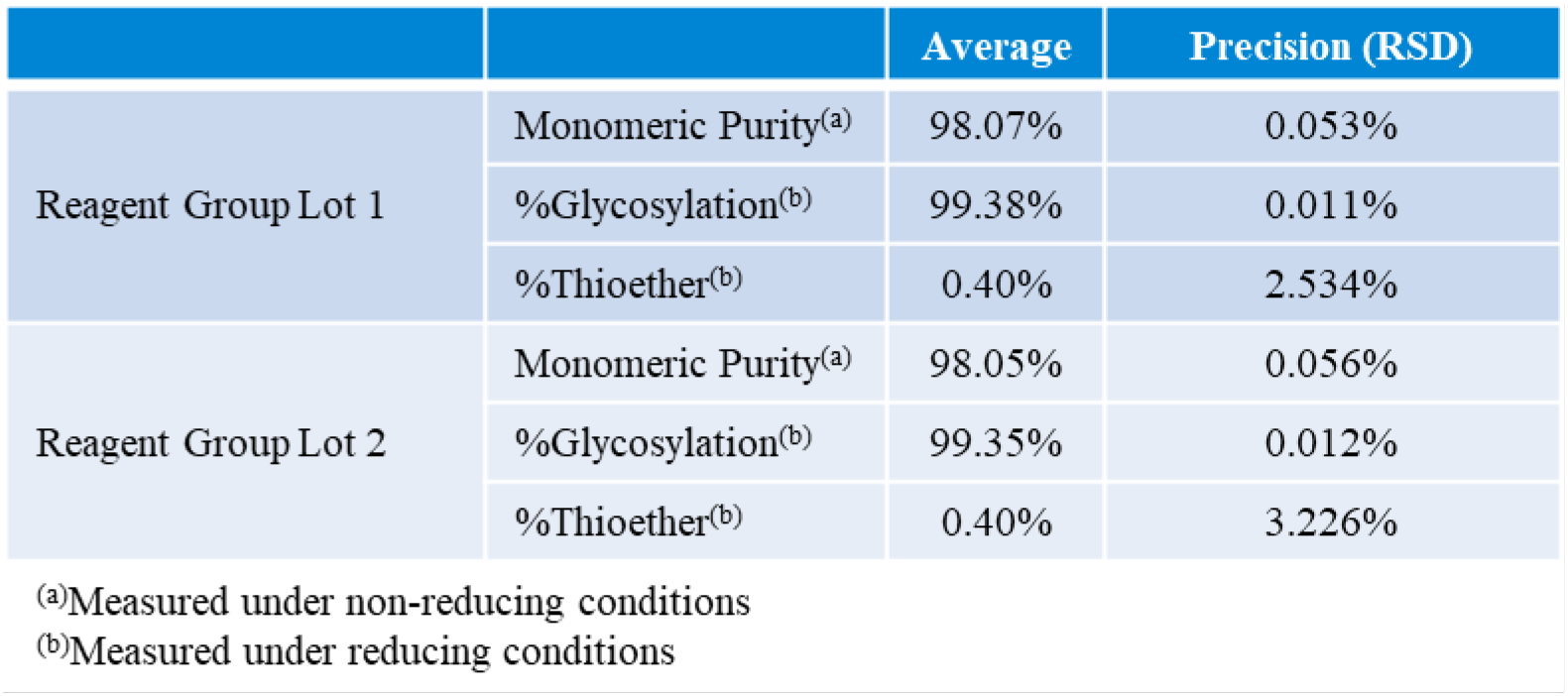
Average monomeric purity, %glycosylation, %thioether, and RSD values for the NISTmAb from two different reagent lot groups used to evaluate the method ruggedness. N = 6.

### Accuracy

The accuracy of the fluorescent CE-SDS method was validated by comparison of the measured results to values collected on a UV CE-SDS system and the published NISTmAb Material Reference Sheet. The values reported by NIST were collected on a traditional single-capillary UV based system [4, 31]. Comparison of reference materials observed values to those theoretically expected is considered an evaluation of method accuracy per ICH guidelines [21]. The monomeric purity value from the ProteoAnalyzer was within 0.5% of the published NISTmAb value, while the percent glycosylation found by the ProteoAnalyzer matched the NISTmAb reference sheet exactly, with both reporting values of 99.38% (Table 6). The percent thioether calculated by the ProteoAnalyzer was within 0.1% of the reported NIST value [22] and consistent with values NIST has reported at different test sites [31].

**Table 6.** Average monomeric purity, %glycosylation, %thioether, and RSD values for the NISTmAb critical quality attributes collected on the fluorescence-based Agilent ProteoAnalyzer system (N = 6) compared to UV based CE-SDS (N = 4) and the NIST-published values collected on a single capillary UV system.

|  | ProteoAnalyzer |  | UV CE-SDS |  | NISTmAb Material Reference Sheet |  |
| --- | --- | --- | --- | --- | --- | --- |
|  | Average | Precision (RSD) | Average | Precision (RSD) | Size Heterogeneity | Expanded Uncertainty, *U* |
| Monomeric Purity<sup>(a)</sup> | 98.07% | 0.053% | 97.60% | 0.137% | 98.49% | 2.19% |
| %Glycosylation<sup>(b)</sup> | 99.38% | 0.011% | 99.37% | 0.015% | 99.38% | 0.09% |
| %Thioether<sup>(b)</sup> | 0.40% | 2.534% | 0.40% | 1.540% | 0.30% | 0.22% |
| <sup>(a)</sup>Measured under non-reducing conditions |  |  |  |  |  |  |
| <sup>(b)</sup>Measured under reducing conditions |  |  |  |  |  |  |

Additionally, the %glycosylation and %thioether were nearly identical between those experimentally measured by fluorescence and UV based CE-SDS. The fluorescent and UV based CE-SDS monomeric purity value experimentally determined had a percent difference of ∼1%. The consistency of the fluorescent CE-SDS method to the experimentally determined UV based CE-SDS and those published by NIST using an orthogonal CE-SDS technique (single capillary UV detection) demonstrate the accuracy of the fluorescent method [21].

### Linearity, LOD, and LOQ

Per ICH Q2R2 guidelines, linearity can be evaluated with a plot of signals as a function of analyte concentration to demonstrate that the analytical procedure is capable of obtaining values that are proportional to the true sample samples across a given range. Test results are evaluated by an appropriate statistical method such as regression line by method of least squares with the correlation coefficient, y-intercept, and slope of the regression line provided [21]. To validate linearity of the fluorescence-based method, the NISTmAb was diluted to ∼1,000, 1,200, 1,500, 1,800, and 2,000 ng/µL in PBS representing a 1,000 ng/µL range and quantified by UV absorption. Three replicates of each concentration were analyzed on the ProteoAnalyzer. For the non-reduced analysis, the average relative concentration (ng/µL) of the intact mAb reported by the ProSize data analysis software from the three replicates was plotted against the absolute concentration as determined by NanoDrop (Figure 2a). For the reduced sample, the relative concentration (ng/µL) reported by the ProSize data analysis software for the L, NGH, H, and thioether was summed to find the total, followed by averaging of the three replicates.

**Fig. 2.**
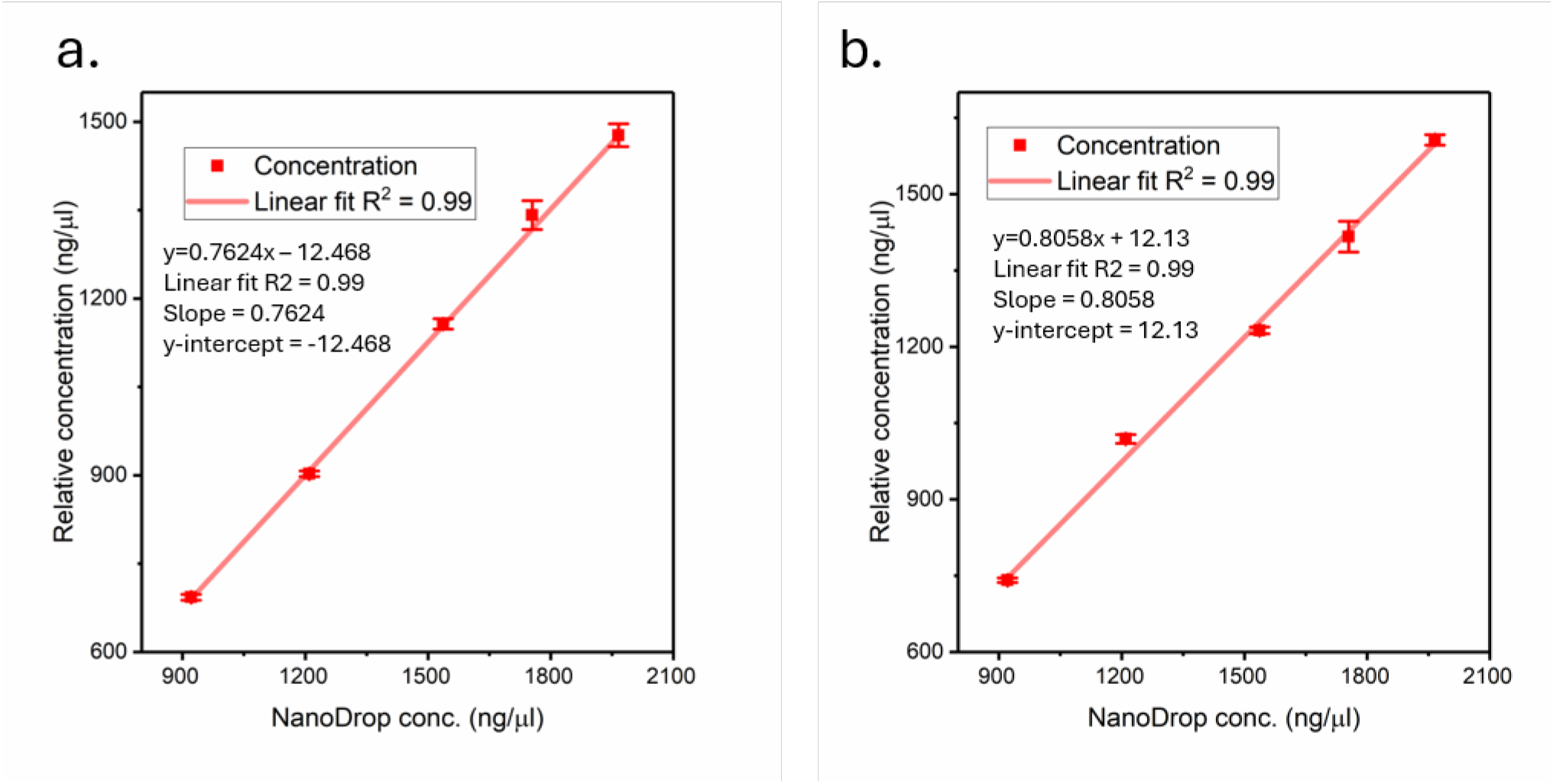
Linearity graphs for the relative concentration determined by the Agilent ProteoAnalyzer system of non-reduced intact NISTmAb (a) and the sum of the reduced L, NGH, H, and thioether (b) against the absolute concentration as determined by NanoDrop. Data shown as the average (N = 3) of the relative concentration ± SD.

The average relative concentration from the ProteoAnalyzer was plotted against the absolute concentration as determined by NanoDrop (Figure 2b). Both the non-reduced and reduced dilution series demonstrated a high degree of linearity with R^2^ values >0.99. The mAb CQAs were analyzed to determine the consistency across the tested range (Tables S6 through S8). The average monomeric purity across the concentration range was 98.14% with a RSD of 0.064%, the percent glycosylation was 99.4% with a RSD of 0.059%, and a percent thioether value of 0.40% with a RSD of 3.434%.

The ProteoAnalyzer has a manufacturer specified linearity quantitative dynamic range of 2 – 2,000 ng/µL for Bovine Serum Albumin (BSA) when diluted in PBS and using a 7 kV for 10 s injection. The LOD and LOQ of the LED fluorescent-based CE-SDS for the NISTmAb was determined through linearity experiments at the bottom of this linear dynamic range (Figure S4). Noise was calculated from a baseline window centered on the mean migration time of the monomer peak and extending ±3 times the peak’s full width at half maximum (FWHM). The LOD and LOQ were determined by interpolating signals corresponding to 3.3-fold and 10-fold the measured noise, respectively, onto the calibration curve [29]. The LOD was found to be 4.16 ± 0.22 ng/µL while the LOQ was found to be 5.02 ± 0.24 ng/µL. The slightly higher LOD/LOQ values may be attributed to the lower injection of 7 kV for 6 s recommended by the manufacturer for non-reduced antibody analysis [23] or due to protein sequence differences between the NISTmAb and BSA.

## Discussion

Therapeutic antibodies as a class are composed of diverse molecules including monoclonal antibodies, bispecific antibodies, antibody drug conjugates, and antibody fragments such as VHH/nanobodies [2]. In this study, the Agilent ProteoAnalyzer system was used to validate an LED fluorescence-based parallel capillary gel electrophoresis method for the non-reduced and reduced analysis of monoclonal antibodies. The validation completed here relied on the use of the NISTmAb as a representative sample. This sample was chosen due to its highly characterized nature [4, 31] allowing for accuracy comparisons to reported values and UV based CE-SDS values experimentally determined [21]. Different therapeutic antibodies may may vary in how close LED fluorescent and UV absorbance CE-SDS values compare due to the differing detection methods. The degree to which robustness/intermediate precision should be evaluated depends on the conditions under which the analytical procedure is expected to be used [21]. In this study, robustness was evaluated with respect to different users and reagent lots. Individual labs should determine what additional variables, if any, should be evaluated.

The majority of the study was focused on the precision of parallel LED fluorescence-based CE-SDS with respect to different experimental variables. We would expect these precision values to be consistent across therapeutic antibody types assuming similar instrumentation and sample preparation was used, though this would need to be experimentally determined for each specific mAb. The experiments presented here can be used as a framework for validating parallel fluorescent-based CE-SDS methods.

The results demonstrated that LED fluorescence-based detection allows for highly precise measurements for the critical quality attributes of monomeric purity, percent glycosylation, and percent thioether. All tests returned RSD values <1% for the percent purity and percent glycosylation CQAs, and <3% for the percent thioether CQA. The higher RSD for the thioether content results from quantitation of a low-concentration impurity and is consistent with previously published CE data [32].

The size and relative concentration of the intact NISTmAb under non-reduced conditions and the L, NGH, H, and thioether fragments under reduced conditions were also analyzed. The reported size for all fragments had precision values of <5% RSD in all tests, with most values <2% RSD. Similar results were obtained for the relative concentrations with all but two being <5% RSD and most having a value <2% RSD. The labeling temperature and 10x Labeling Buffer volume robustness tests showed slightly higher RSD values for the NGH relative concentration at 5.728% and 8.653% respectively, presumably resulting from the low level of this impurity [32]. Despite the slightly higher variability in the NGH relative concentrations, the percent glycosylation and percent thioether CQAs for the labeling temperature (0.037% and 3.867% respectively) and the 10x Labeling Buffer volume (0.038% and 2.839% respectively) remained consistent, indicating that this had minimal impact on the CQA variability.

In addition to the high reproducibility of the LED fluorescence-based method, the stable baseline achieved with fluorescent CE-SDS allows for greater confidence in the specificity of the assay. The well-characterized baseline drift found in UV (Figure 1 and S2) increase the likelihood of miscalling or mis quantifying low-level impurities [12, 14-16] and usually requires the subtraction of a “blank” to increase confidence in peak calling [11]. This can be challenging due to injection-to-injection baseline drift differences (Figure S2). For instance, two of the UV CE-SDS baselines had a positive slope throughout the run while the other two demonstrated a negative sloping baseline. In contrast, blank subtraction was not required for the fluorescent method, while still maintaining high precision results due to the baseline stability seen in all injections (Figures 1 and S1).

While fluorescence-based assays bring advantages to monoclonal antibody analysis, most historical data has been collected using UV absorption-based systems. To validate the LED fluorescence-based method, the CQAs were compared to traditional UV absorption values collected during this study and published by the US National Institute of Standards (NIST) [22]. All results obtained in this study were consistent with those reported in the NIST Material Reference Data Sheet, indicating that LED fluorescence-based methods can be used for CQA analysis of therapeutic mAbs.

## Conclusion

With the growing demand for therapeutic antibodies, higher-throughput methodologies with more stable baselines are required. Traditionally, CE-SDS has been conducted on single capillary systems which limit sample throughput. Common electrophoresis times on these single capillary systems are around 20 min per sample and are able to analyze up to 96 samples without user intervention. In contrast, the fluorescent based system used in this study allows for parallel analysis of 12 samples in around 30 min and can analyze up to 288 samples without user intervention. While fluorescent based higher-throughput alternatives are available [33] published and validated CE-SDS methods by groups such as the USP rely on UV absorption for detection of sample, which limits the adoption of fluorescent methods [8]. The results from this study demonstrate the high specificity, robustness, ruggedness, accuracy, and linearity of LED fluorescence based parallel capillary gel electrophoresis method. The data presented here provides a starting point for the validation of individual therapeutic monoclonal antibodies.

## Author contributions

KDL and MG contributed to the study conception and design. Fluorescent CE-SDS investigation and data curation was completed by KDL and WAP. UV CE-SDS investigation and data curation was completed by SYH and KDL. The original draft of the manuscript was written by KDL and all authors reviewed/edited. All authors read and approved the final manuscript.

## References

1. Therapeutic monoclonal antibodies approved or in regulatory review [Internet]. Available from: https://www.antibodysociety.org/antibody-therapeutics-product-data.

2. Chiu ML, Goulet DR, Teplyakov A, Gilliland GL. Antibody Structure and Function: The Basis for Engineering Therapeutics. Antibodies. 2019;8(4):55. 10.3390/antib8040055

3. Kaur H. Chapter 2 - Physicochemical characterization of monoclonal antibodies. In: Kaur H, Reusch D, editors. Monoclonal Antibodies: Academic Press; 2021. p. 31–63. 10.1016/B978-0-12-822318-5.00007-7

4. Turner A, Yandrofski K, Telikepalli S, King J, Heckert A, Filliben J, et al. Development of orthogonal NISTmAb size heterogeneity control methods. Anal Bioanal Chem. 2018;410(8):2095–110. 10.1007/s00216-017-0819-3

5. Zhang Q, Schenauer MR, McCarter JD, Flynn GC. IgG1 thioether bond formation in vivo. J Biol Chem. 2013;288(23):16371–82. 10.1074/jbc.M113.468397

6. Tous GI, Wei Z, Feng J, Bilbulian S, Bowen S, Smith J, et al. Characterization of a Novel Modification to Monoclonal Antibodies: Thioether Cross-link of Heavy and Light Chains. Anal Chem. 2005;77(9):2675–82. 10.1021/ac0500582

7. Song X, Tian H, Jing R, Liu K, Xu K, Guo L, et al. Navigating the Frontier: Advances in monoclonal antibody purity control. Protein Expression Purif. 2025;232:106725. 10.1016/j.pep.2025.106725

8. General Chapter <129> Analytical Procedures for Recombinant Therapeutic Monoclonal Antibodies. United States Pharmacopeia and National Formulary (USP-NF) 2025. 10.31003/USPNF_M6297_03_01

9. Kumar R, Guttman A, Rathore AS. Applications of capillary electrophoresis for biopharmaceutical product characterization. Electrophoresis. 2022;43(1-2):143–66. 10.1002/elps.202100182

10. Bello MS, de Besi P, Righetti PG. Thermally induced fluctuations of the electric current and baseline in capillary electrophoresis. J Chromatogr A. 1993;652(2):317–27. 10.1016/0021-9673(93)83249-R

11. Gu Y, Voronov S, Ding J, Mussa N, Li ZJ. Assessment of CE-based baseline disturbances using simulation and targeted experimental evaluation-impact on the purity determination of therapeutic proteins. Anal Bioanal Chem. 2019;411(11):2425–37. 10.1007/s00216-019-01704-6

12. Xu X, Kok WT, Poppe H. Noise and baseline disturbances in indirect UV detection in capillary electrophoresis. J Chromatogr A. 1997;786(2):333–45. 10.1016/S0021-9673(97)00616-X

13. Štěpánová S, Kašička V. Applications of capillary electromigration methods for separation and analysis of proteins (2017–mid 2021) – A review. Analytica Chimica Acta. 2022;1209:339447. 10.1016/j.aca.2022.339447

14. Luo J, Sänger-van de Griend C, editors. Troubleshooting CE-SDS: baseline disturbances, peak area repeatability and the presensence of ghost peaks. CASSS CE Pharm; 2015.https://www.casss.org/docs/default-source/ce-pharm/reports-troubleshooting-workshops/troubleshooting-ce-sds---baseline-disturbances---peak-area-repeatability-and-the-presence-of-ghost-peaks.pdf

15. Rodat-Boutonnet A, Naccache P, Morin A, Fabre J, Feurer B, Couderc F. A comparative study of LED-induced fluorescence and laser-induced fluorescence in SDS-CGE: application to the analysis of antibodies. Electrophoresis. 2012;33(12):1709–14. 10.1002/elps.201200132

16. Zhang Z, Park J, Barrett H, Dooley S, Davies C, Verhagen MF. Capillary Electrophoresis-Sodium Dodecyl Sulfate with Laser-Induced Fluorescence Detection as a Highly Sensitive and Quality Control-Friendly Method for Monitoring Adeno-Associated Virus Capsid Protein Purity. Hum Gene Ther. 2021;32(11-12):628–37. 10.1089/hum.2020.233

17. Michels DA, Brady LJ, Guo A, Balland A. Fluorescent Derivatization Method of Proteins for Characterization by Capillary Electrophoresis-Sodium Dodecyl Sulfate with Laser-Induced Fluorescence Detection. Anal Chem. 2007;79(15):5963–71 10.1021/ac0705521

18. Gennaro L, Felten C, Salas-Salano O. Optimization Approaches in the Routine Analysis of Monoclonal Anitbodies by Capillary Electrophoresis. Optimization. 21(12).

19. Griffin S. Fused Silica Capillary: the Story Behind the Technology. LC GC North America. 2002;20:928.

20. Altria KD. Improved Performance in Capillary Electrophoresis using Internal Standards. LC GC Europe. 2002;15:588–94.

21. Use ICfHoTRfPfH. Q2(R2) Validation of Analytical Procedures: Guidance for Industry. US Food and Drug Administration; 2024.

22. 2022 SRM 8671; NISTmAb, Humanized IgG1κ Monoclonal Antibody Reference Material Information Sheet. Gaithersburg, MD: United States National Institute of Standards and Technology; 24 January 2022. Report No.: Lot 14HB-D-002.

23. Analysis of NIST Antibody on the Agilent ProteoAnalyzer System. Agilent Technologies technical overview. 2024. 5994–6781EN

24. Liu Q, Hong J, Zhang Y, Wang Q, Xia Q, Knierman MD, et al. Rapid identification of antibody impurities in size-based electrophoresis via CZE-MS generated spectral library. Sci Rep. 2024;14(1):20239. 10.1038/s41598-024-70914-5

25. Luttgeharm K, Pike W. Accurate mAb Sizing Using the NIST mAb as a Ladder for the Agilent ProteoAnalyzer System. 2025. 5994–8815EN. Available from: https://www.agilent.com/cs/library/applications/an-antibody-sizing-nist-proteoanalyzer-5994-8815en-agilent.pdf.

26. Bean SR, Lookhart GL. Sodium Dodecyl Sulfate Capillary Electrophoresis of Wheat Proteins. 1. Uncoated Capillaries. J Agric Food Chem. 1999;47(10):4246–55. 10.1021/jf990413n

27. Guttman A, Filep C, Karger BL. Fundamentals of Capillary Electrophoretic Migration and Separation of SDS Proteins in Borate Cross-Linked Dextran Gels. Anal Chem. 2021;93(26):9267–76. 10.1021/acs.analchem.1c01636

28. Noise Determination. Agilent OpenLAB CDS: Agilent Technologies; 2016.

29. Armbruster DA, Pry T. Limit of blank, limit of detection and limit of quantitation. Clin Biochem Rev. 2008;29 Suppl 1(Suppl 1):S49–52.

30. Sanger-van de Griend CE. CE-SDS method development, validation, and best practice-An overview. Electrophoresis. 2019;40(18-19):2361–74. 10.1002/elps.201900094

31. Schiel JE, Turner A, Mouchahoir T, Yandrofski K, Telikepalli S, King J, et al. The NISTmAb Reference Material 8671 value assignment, homogeneity, and stability. Anal Bioanal Chem. 2018;410(8):2127–39. 10.1007/s00216-017-0800-1

32. Luraschi P, Infusino I, Merlotti C, Franzini C. Analytical variation in the measurement of serum monoclonal component by capillary electrophoresis. Clin Chim Acta. 2004;349(1-2):151–6. 10.1016/j.cccn.2004.06.016

33. Kahle J, Maul KJ, Wätzig H. The next generation of capillary electrophoresis instruments: Performance of CE-SDS protein analysis. Electrophoresis. 2018;39(2):311–25. 10.1002/elps.201700278

