## Supplementary material for "Validation of a high-throughput fluorescent Capillary Electrophoresis Sodium Dodecyl Sulfate method for monoclonal antibody size heterogeneity assessment": Supplmental Information

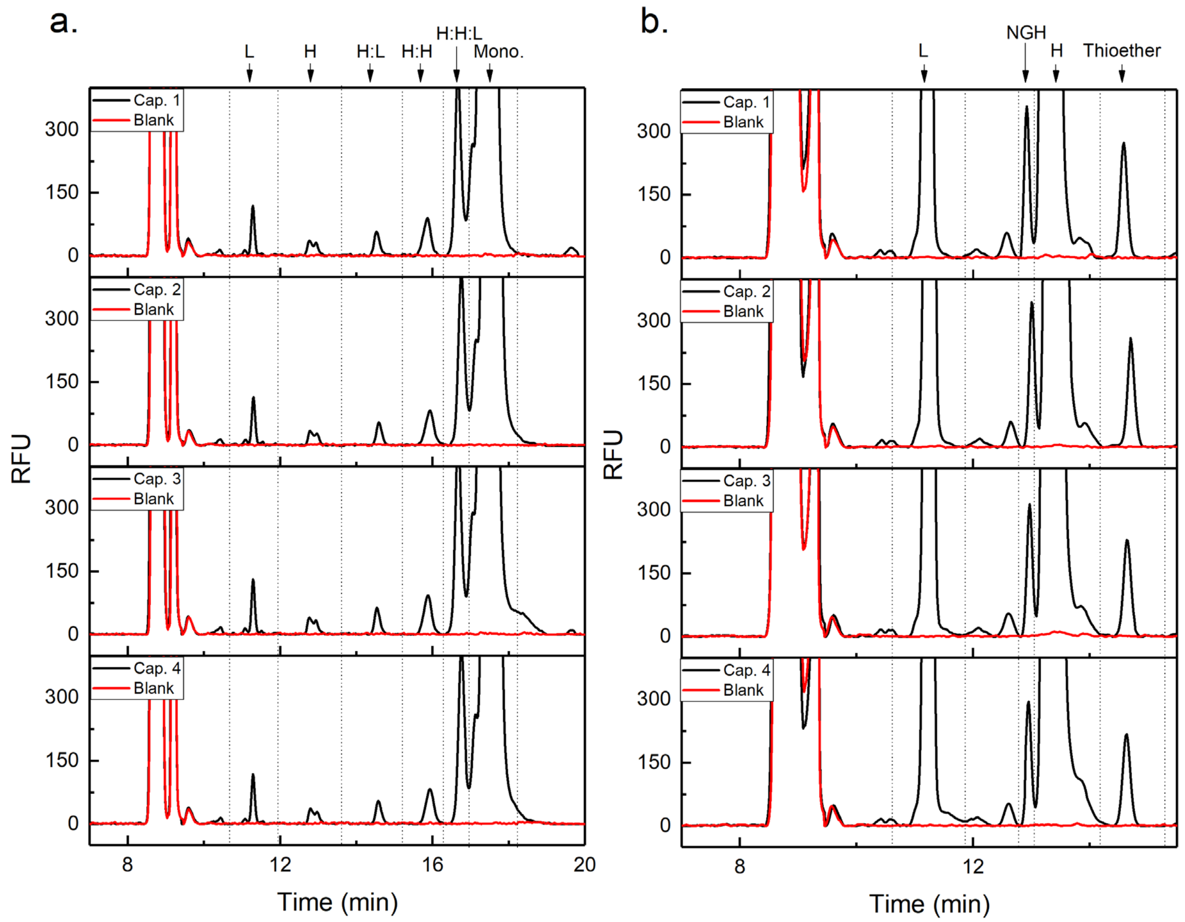


**Fig. S1** Representative electropherograms of the (a) non-reduced NISTmAb and blank from 4 capillaries using the LED-induced fluorescence-based method for monomeric purity and (b) reduced NISTmAb peak with blank run for percent glycosylation and thioether. L = Light Chain, NGH = Non-glycosylated Heavy Chain, H = Heavy Chain, Mono = Monomer.

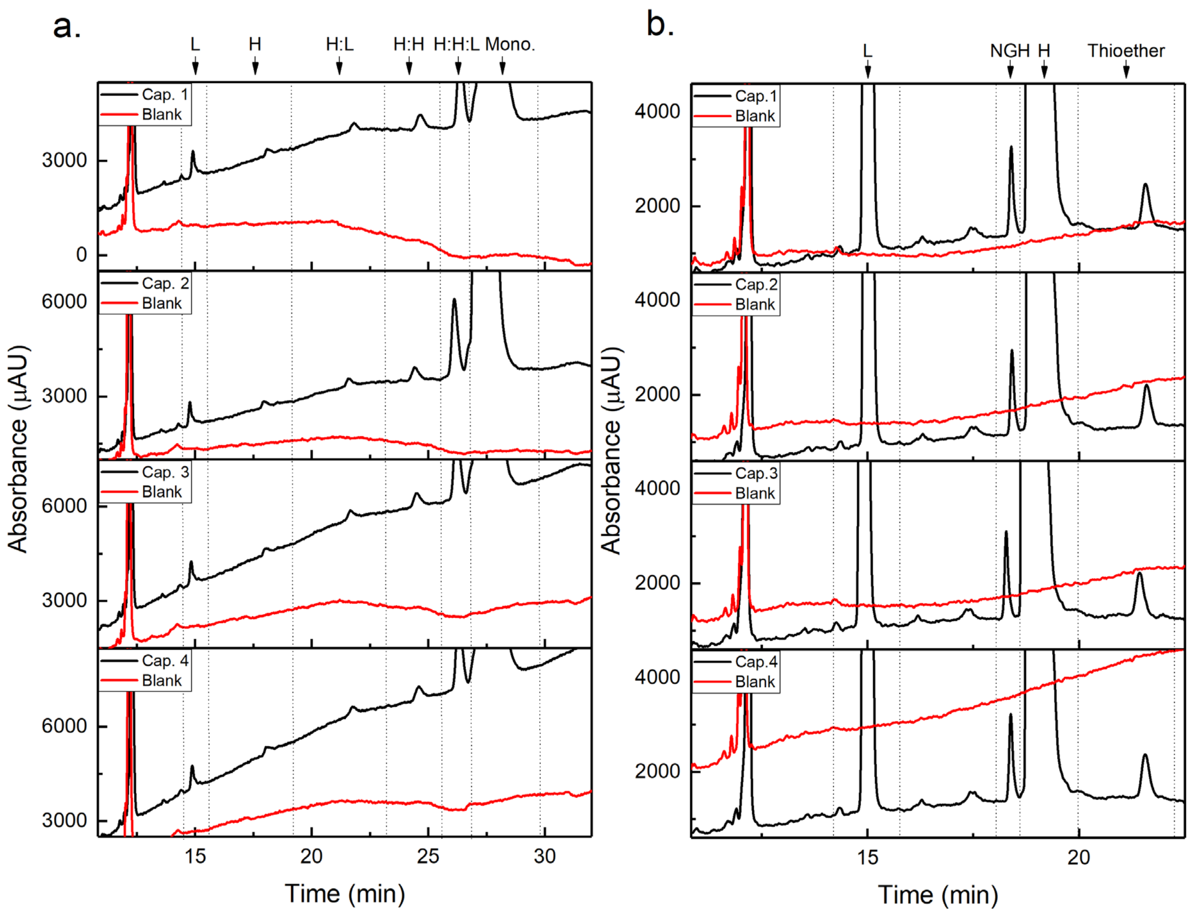


**Fig. S2** Representative electropherograms of the (a) non-reduced NISTmAb and blank from 4 using the LED-induced fluorescence-based method and (b) reduced NISTmAb, peak with blank run for percent glycosylation and thioether. L = Light Chain, NGH = Non-glycosylated Heavy Chain, H = Heavy Chain, Mono = Monomer.

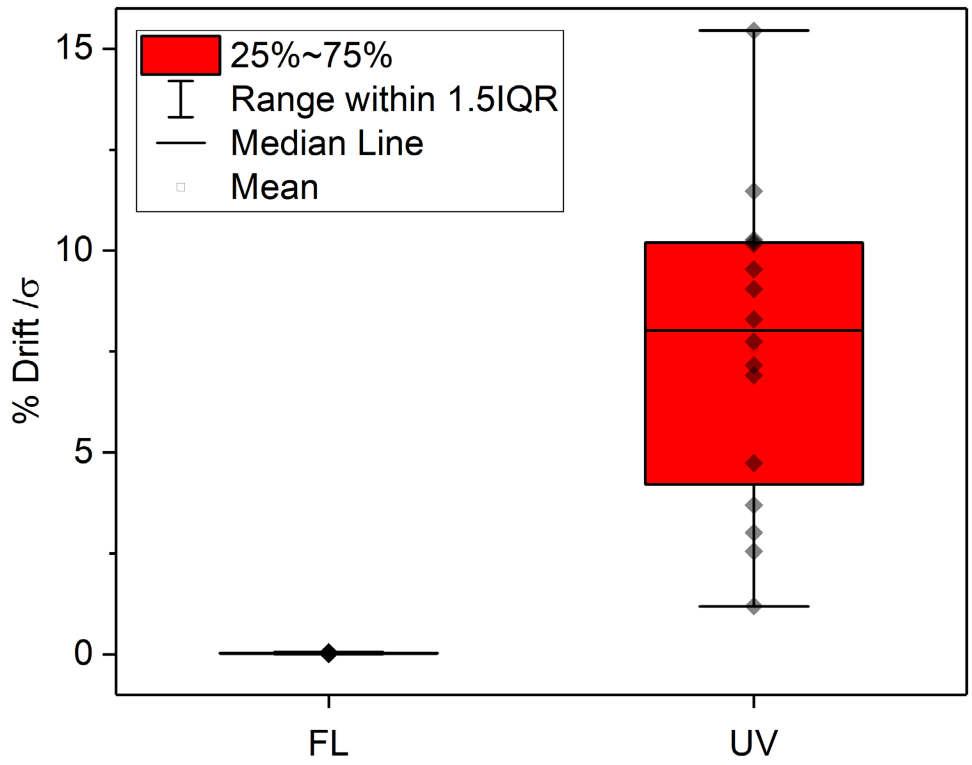


**Fig. S3** Baseline drift analysis of the led-based fluorescence (FL) detector on the ProteoAnalyzer and the UV detector on the BioPhase 8800. Each data point represents one of the % Drift /$\sigma$ windows analyzed. The data was analyzed via a box and whisker plot (FL n = 20, UV n =16). The much larger drift-to-noise values observed on the BioPhase 8800 indicate a less stable baseline.

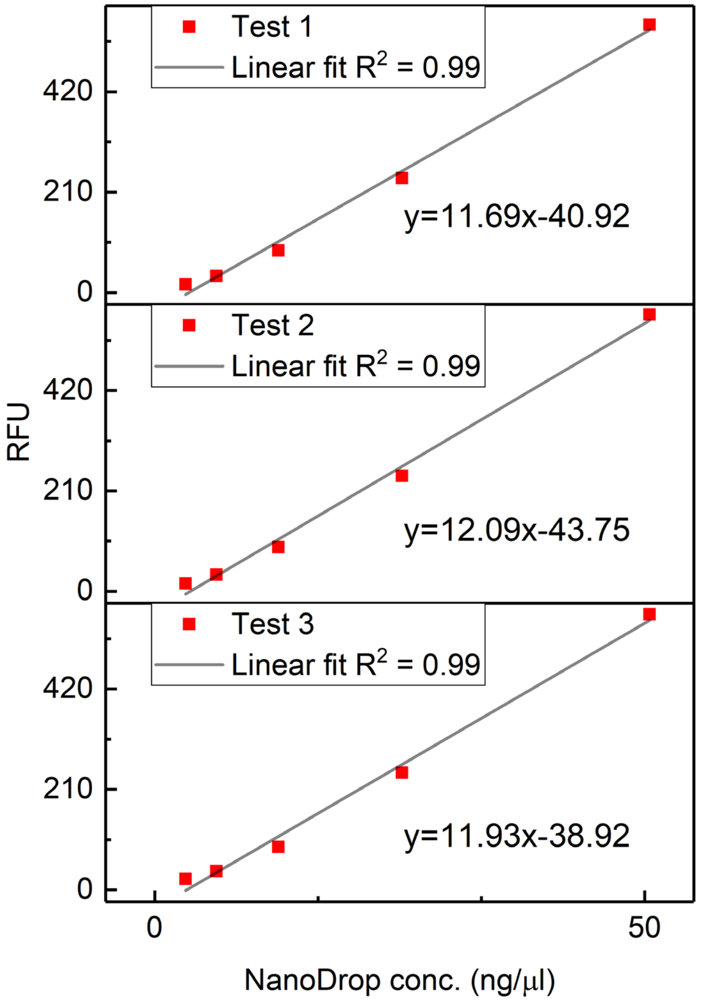


**Fig S4.** Calibration curves used to calculate the limit of detection (LOD) and limit of quantification (LOQ) for the fluorescent CE-SDS method. All curves showed an $R^2$ value of >0.99.

**Table S1.** Size (kDa) and relative concentration (ng/µL) for the intact monomer under non-reduced conditions and the L, NGH, H, and thioether under reduced conditions. Data are the average of six independent capillaries used to evaluate the precision of the LED fluorescence-based parallel CE instrument. N = 6. L = Light chain, NGH = Non-glycosylated heavy chain, H = Heavy chain

| | Size (kDa) | | Concentration (ng/µL) | |
|---|---|---|---|---|
| | Average | Precision (RSD) | Average | Precision (RSD) |
| Monomer[(a)] | 147.82 | 1.311% | 1134.70 | 0.831% |
| L[(b)] | 26.95 | 1.380% | 337.73 | 1.413% |
| NGH[(b)] | 56.9 | 1.990% | 5.19 | 2.602% |
| H[(b)] | 63.65 | 3.000% | 829.45 | 1.402% |
| Thioether[(b)] | 111.43 | 2.990% | 4.71 | 2.814% |

[(a)]Measured under non-reducing conditions

[(b)]Measured under reducing conditions

**Table S2.** Size (kDa) and relative concentration (ng/µL) for the intact monomer under non-reduced conditions and the L, NGH, H, and thioether under reduced conditions. Data are the average of six samples individually mixed 1:29 (sample : Labeling Buffer mix) to evaluate the method precision of the LED fluorescence-based parallel CE protocol. The higher RSDs observed for the NGH and Thioether concentrations are presumably due to the low level of these impurities combined with the individual mixing of sample and labeling buffer mix. N = 6. L = Light chain, NGH = Non-glycosylated heavy chain, H = Heavy chain

| | Size (kDa) | | Concentration (ng/µL) | |
|---|---|---|---|---|
| | Average | Precision (RSD) | Average | Precision (RSD) |
| Monomer[(a)] | 142.82 | 1.551% | 1012.66 | 3.793% |
| L[(b)] | 26.90 | 1.051% | 328.81 | 2.819% |
| NGH[(b)] | 56.45 | 1.914% | 5.17 | 4.935% |
| H[(b)] | 62.82 | 2.544% | 823.90 | 2.929% |
| Thioether[(b)] | 109.60 | 2.459% | 4.74 | 3.496% |

[(a)]Measured under non-reducing conditions

[b)]Measured under reducing conditions

**Table S3.** Size (kDa) and relative concentration (ng/µL) for the intact monomer under non-reduced conditions and the L, NGH, H, and thioether under reduced conditions for the values used to evaluate the method robustness of the LED fluorescence-based parallel CE protocol. N = 3. L = Light chain, NGH = Non-glycosylated heavy chain, H = Heavy chain

| | | Size (kDa) | | Concentration (ng/µL) | |
|---|---|---|---|---|---|
| | | Average | Precision (RSD) | Average | Precision (RSD) |
| Injection Voltage | Monomer[a] | 148.44 | 1.992% | 1138.04 | 0.799% |
| | L[b] | 26.97 | 1.729% | 338.42 | 1.070% |
| | NGH[b] | 56.66 | 2.774% | 5.28 | 2.743% |
| | H[b] | 63.46 | 3.976% | 833.47 | 1.392% |
| | Thioether[b] | 110.60 | 4.021% | 4.73 | 1.633% |
| Electrophoresis Voltage | Monomer[a] | 147.82 | 1.196% | 1135.25 | 1.079% |
| | L[b] | 27.20 | 1.471% | 330.26 | 1.927% |
| | NGH[b] | 57.54 | 2.294% | 5.13 | 3.114% |
| | H[b] | 64.81 | 3.683% | 815.25 | 1.511% |
| | Thioether[b] | 112.97 | 3.535% | 4.67 | 2.047% |
| Labeling Temperature | Monomer[a] | 143.44 | 3.623% | 1163.43 | 2.797% |
| | L[b] | 27.23 | 1.818% | 393.33 | 3.273% |
| | NGH[b] | 57.44 | 2.743% | 6.19 | 5.728% |
| | H[b] | 64.80 | 4.257% | 962.34 | 1.107% |
| | Thioether[b] | 112.90 | 4.094% | 5.46 | 3.786% |
| Labeling Buffer Volume | Monomer[a] | 141.78 | 4.262% | 1148.68 | 2.535% |
| | L[b] | 27.14 | 1.416% | 386.65 | 3.187% |
| | NGH[b] | 57.41 | 2.004% | 5.96 | 8.653% |
| | H[b] | 64.52 | 3.314% | 933.54 | 3.182% |
| | Thioether[b] | 113.09 | 2.826% | 5.28 | 3.246% |

[a]Measured under non-reducing conditions

[b]Measured under reducing conditions

**Table S4.** Size (kDa) and relative concentration (ng/µL) for the intact monomer under non-reduced conditions and the L, NGH, H, and thioether under reduced conditions from two different users used to evaluate the method ruggedness of the LED fluorescence-based parallel CE protocol. N = 6. L = Light chain, NGH = Non-glycosylated heavy chain, H = Heavy chain

| | | Size (kDa) | | Concentration (ng/µL) | |
|---|---|---|---|---|---|
| | | Average | Precision (RSD) | Average | Precision (RSD) |
| User 1 | Monomer[a] | 144.78 | 3.535% | 1228.68 | 1.197% |
| | L[b] | 26.37 | 1.167% | 345.03 | 0.789% |
| | NGH[b] | 55.40 | 1.823% | 5.44 | 2.150% |
| | H[b] | 61.45 | 1.536% | 829.97 | 1.053% |
| | Thioether[b] | 107.30 | 2.816% | 4.77 | 2.911% |
| User 2 | Monomer[a] | 136.42 | 1.577% | 1095.79 | 1.787% |
| | L[b] | 26.73 | 1.373% | 333.79 | 0.717% |
| | NGH[b] | 56.53 | 2.165% | 5.31 | 1.472% |
| | H[b] | 62.93 | 2.746% | 820.93 | 1.534% |
| | Thioether[b] | 110.53 | 3.107% | 4.57 | 1.743% |

[a] Measured under non-reducing conditions

[b] Measured under reducing conditions

**Table S5.** Size (kDa) and relative concentration (ng/µL) for the intact monomer under non-reduced conditions and the L, NGH, H, and Thioether under reduced conditions from two different users used to evaluate the method ruggedness of the LED fluorescence-based parallel CE protocol. N = 6. L = Light chain, NGH = Non-glycosylated heavy chain, H = Heavy chain

| | | Size (kDa) | | Concentration (ng/µL) | |
|---|---|---|---|---|---|
| | | Average | Precision (RSD) | Average | Precision (RSD) |
| Lot Group 1 | Monomer[(a)] | 147.82 | 1.311% | 1134.70 | 0.831% |
| | L[(b)] | 26.95 | 1.383% | 337.73 | 1.413% |
| | NGH[(b)] | 56.90 | 1.991% | 5.19 | 2.602% |
| | H[(b)] | 63.65 | 2.997% | 829.45 | 1.402% |
| | Thioether[(b)] | 111.43 | 2.986% | 4.71 | 2.814% |
| Lot Group 2 | Monomer[(a)] | 144.78 | 3.535% | 1228.68 | 1.197% |
| | L[(b)] | 26.37 | 1.167% | 345.03 | 0.789% |
| | NGH[(b)] | 55.40 | 1.823% | 5.44 | 2.150% |
| | H[(b)] | 61.45 | 1.536% | 829.97 | 1.053% |
| | Thioether[(b)] | 107.30 | 2.816% | 4.77 | 2.911% |

[(a)]Measured under non-reducing conditions

[(b)]Measured under reducing conditions

**Table S6.** Monomeric purity, size (kDa), relative concentration (ng/µL) as determined by the Agilent ProteoAnalyzer system and NanoDrop concentration used for linearity calculations of the intact monomer under non-reduced conditions. N = 3.

| | Monomeric Purity | | Size (kDa) | | ProteoAnalyzer Relative Concentration (ng/µL) | | |
|---|---|---|---|---|---|---|---|
| Sample | Average | Precision (RSD) | Average | Precision (RSD) | Average | Precision (RSD) | NanoDrop Absolute Concentration (ng/µL) |
| 2000 ng/µL | 98.10% | 0.000% | 147.10 | 0.556% | 1477.20 | 1.308% | 1966 |
| 1800 ng/µL | 98.13% | 0.059% | 148.53 | 0.967% | 1341.78 | 1.825% | 1755 |
| 1500 ng/µL | 98.10% | 0.000% | 146.73 | 0.925% | 1156.88 | 0.755% | 1537 |
| 1200 ng/µL | 98.13% | 0.059% | 147.47 | 1.897% | 902.42 | 0.529% | 1210 |
| 1000 ng/µL | 98.23% | 0.059% | 146.73 | 2.013% | 693.06 | 0.707% | 921 |

**Table S7.** %glycosylation, %thioether, total relative concentration (L+H+NGH+H+thioether ng/µL) as determined by the Agilent ProteoAnalyzer system, and NanoDrop concentration used for linearity calculations of the NISTmAb under reduced conditions. N = 3.

| | % Glycosylation | | % Thioether | | ProteoAnalyzer Relative Concentration (ng/µL) | | |
|---|---|---|---|---|---|---|---|
| Sample | Average | Precision (RSD) | Average | Precision (RSD) | Average | Precision (RSD) | NanoDrop Absolute Concentration (ng/µL) |
| 2000 ng/µL | 99.32% | 0.057% | 0.41% | 1.641% | 1606.57 | 0.629% | 1966 |
| 1800 ng/µL | 99.35% | 0.016% | 0.39% | 2.033% | 1416.23 | 2.130% | 1755 |
| 1500 ng/µL | 99.36% | 0.006% | 0.41% | 3.188% | 1232.19 | 0.551% | 1537 |
| 1200 ng/µL | 99.43% | 0.043% | 0.40% | 5.711% | 1018.79 | 0.830% | 1210 |
| 1000 ng/µL | 99.45% | 0.008% | 0.39% | 3.031% | 741.03 | 0.623% | 921 |

**Table S8.** Average size for the L, NGH, H, and thioether species of the reduced linearity samples as determined by the Agilent ProteoAnalyzer system. N = 3. L = Light chain, NGH = Non-glycosylated heavy chain, H = Heavy chain

| | L | | NGH | | H | | Thioether | |
|---|---|---|---|---|---|---|---|---|
| Sample | Average | Precision (RSD) | Average | Precision (RSD) | Average | Precision (RSD) | Average | Precision (RSD) |
| 2000 ng/µL | 26.80 | 0.746% | 55.70 | 0.54% | 61.47 | 0.470% | 107.30 | 0.646% |
| 1800 ng/µL | 26.83 | 1.721% | 56.03 | 1.75% | 61.73 | 1.470% | 108.37 | 2.611% |
| 1500 ng/µL | 26.70 | 1.350% | 55.33 | 1.64% | 61.13 | 1.484% | 106.50 | 2.662% |
| 1200 ng/µL | 27.03 | 0.214% | 57.00 | 0.61% | 63.23 | 1.076% | 111.00 | 0.624% |
| 1000 ng/µL | 26.97 | 1.302% | 56.70 | 1.86% | 63.57 | 2.970% | 110.57 | 2.759% |
